# Endothelial β1 Integrins are Necessary for Microvascular Function and Glucose Uptake

**DOI:** 10.1101/2024.08.18.607045

**Authors:** Nathan C. Winn, Deborah A. Roby, P. Mason McClatchey, Ian M. Williams, Deanna P. Bracy, Michelle N. Bedenbaugh, Louise Lantier, Erin J. Plosa, Ambra Pozzi, Roy Zent, David H. Wasserman

**Affiliations:** Department of Molecular Physiology & Biophysics, Vanderbilt University, Nashville, TN, USA; Vanderbilt Mouse Metabolic Phenotyping Center, Vanderbilt University, Nashville, TN, USA; Department of Pediatrics, Vanderbilt University Medical Center, Nashville, TN, USA; Department of Medicine, Vanderbilt University Medical Center, Nashville, TN, USA; Veterans Affairs, Nashville, TN, USA

## Abstract

Microvascular insulin delivery to myocytes is rate limiting for the onset of insulin-stimulated muscle glucose uptake. The structural integrity of capillaries of the microvasculature is regulated, in part, by a family of transmembrane adhesion receptors known as integrins, which are composed of an α and β subunit. The integrin β1 (itgβ1) subunit is highly expressed in endothelial cells (EC). EC itgβ1 is necessary for the formation of capillary networks during embryonic during development and its knockdown in adult mice blunts the reactive hyperemia that manifests during ischemia reperfusion. In this study we investigated the contribution of skeletal muscle EC itgβ1 in microcirculatory function and glucose uptake. We hypothesized that loss of EC itgβ1 would impair microvascular hemodynamics and glucose uptake during insulin stimulation, creating ‘delivery’-mediated insulin resistance. An itgβ1 knockdown mouse model was developed to avoid lethality of embryonic gene knockout and the deteriorating health resulting from early post-natal inducible gene deletion. We found that mice with (itgβ1^fl/fl^SCLcre) and without (itgβ1^fl/fl^) inducible stem cell leukemia cre recombinase (SLCcre) expression at 10 days post cre induction have comparable exercise tolerance and pulmonary and cardiac functions. We quantified microcirculatory hemodynamics using intravital microscopy and the ability of mice to respond to the high metabolic demands of insulin-stimulated muscle using a hyperinsulinemic-euglycemia clamp. We show that itgβ1^fl/fl^SCLcre mice compared to itgβ1^fl/fl^ littermates have, i) deficits in capillary flow rate, flow heterogeneity, and capillary density; ii) impaired insulin-stimulated glucose uptake despite sufficient transcapillary insulin efflux; and iii) reduced insulin-stimulated glucose uptake due to perfusion-limited glucose delivery. Thus, EC itgβ1 is necessary for microcirculatory function and to meet the metabolic challenge of insulin stimulation.

## Introduction

Decreased tissue glucose uptake in response to insulin is a central feature of obesity and diabetes (1). Muscle glucose uptake accounts for approximately 85% of whole-body glucose uptake during a hyperinsulinemic-euglycemic clamp (1). As the bulk of insulin sensitive tissue, uptake of glucose by skeletal muscle is essential to an understanding of whole body glucoregulation. Indeed, deficits in muscle glucose uptake can directly cause impaired glucose tolerance, hyperglycemia, and is a primary feature in the development of type 2 diabetes. Muscle glucose uptake requires that glucose be delivered to the muscle membrane by the vascular system, transported across the membrane, and phosphorylated within the cells. This ‘distributed control’ is how blood glucose can be maintained within a narrow range even during high metabolic demands of insulin stimulation or exercise (2). The extra-myocellular milieu represents an important site of resistance to muscle glucose uptake. This milieu includes i) extracellular matrix (ECM) components (3) which interact with a family of cell adhesion receptors known as integrins and ii) the microcirculation (4). Microcirculatory blood flow, capillary surface area, and the structural integrity of capillaries determine the delivery of perfusion-limited molecules (e.g., glucose, O_2_) and dispersion-limited molecules (e.g., insulin) to tissues. In contrast to glucose, which is tightly regulated by capillary blood flow, insulin dispersion from capillaries to the interstitial space is determined by the permeability of capillaries to insulin and number of perfused capillaries that provide endothelial cell (EC) surface area for exchange (5, 6). Importantly, microvascular insulin delivery to myocytes is rate limiting for the onset of insulin-stimulated muscle glucose uptake (7–11). This is clinically relevant given that patients with type 2 diabetes have reduced capillary density and deficits in perfusion in skeletal muscle (12, 13). These deficits are also associated with increases in muscle collagens and other ECM components that are directly linked to insulin resistance (14–16). Together, these data suggest that microvascular dysfunction and capillary rarefaction are mechanisms by which ECM remodeling may mediate muscle insulin resistance. Understanding the molecular and physiological mechanisms by which ECM-integrin interactions affect EC function may lead to the identification of new therapeutic targets that enhance microvascular function and insulin sensitivity.

The structural integrity and formation of capillaries is regulated, in part, by integrins (17). These transmembrane receptors are composed of α and β heterodimers that modulate cell signaling, proliferation, migration, differentiation, and metabolism (18, 19). In mammals, there are 18 α (itgα) and 8 β subunits (itgβ) that combine to form 24 distinct receptors. Integrins containing the β1 subunit (itgβ1) are abundant in ECs of the microvasculature (19). Specifically, EC itgβ1 is necessary for the formation of capillary networks during embryonic development (20–24) and is required to maintain cell-cell junction integrity by regulating the localization of VE-cadherin (20, 22). In addition to its role in regulating EC function during development and cell junction integrity (20, 22, 25, 26), itgβ1 is involved in sensing and responding to blood flow (27). *In vitro* studies show that vascular shear forces are sensed by itgβ1 which signal increased expression of endothelial nitric oxide synthase (eNOS) (28). Activation of eNOS generates nitric oxide, which causes activation of protein kinase G (PKG). PKG causes smooth muscle relaxation and increases angiogenesis (29, 30). Administration of itgβ1 blocking antibodies or deletion of itgβ1 in ECs in 10-week-old mice using the VE-cadherin cre mouse inhibits flow-mediated dilation through an eNOS-specific mechanism in a mouse hindlimb ischemia model (25). Despite seminal work defining fundamental EC itgβ1 functions in vascular growth and regulation *in vitro* (19), the physiological role of this integrin subunit in regulating glucose uptake and insulin access to target tissues in vivo is not known. This is important to know because the integrated nature of insulin action requires study of the whole organism in which the vascular and muscle system is intact.

As mentioned above, the ability of insulin to access skeletal muscle is dependent on the rate of blood flow, the capillary surface area for exchange and the permeability of those capillaries to insulin. Hemodynamic factors that control blood flow and capillary recruitment are upstream mechanisms for insulin delivery that have been studied in detail (31). In contrast, the permeability of capillaries to insulin is a distinctly local factor. The physiology of insulin egress from the microvasculature to the interstitium has been difficult to examine in vivo. However, an in vivo intravital microscopy technique has made it possible to measure trans-endothelial efflux of insulin using a bioactive B1-Alexa Fluor 647-insulin conjugate (5, 6, 32). In this study, we leverage this novel technique to understand the extent to which EC integrins control microvascular hemodynamics and trans-endothelial insulin efflux in skeletal muscle. We specifically studied skeletal muscle because it is 1) densely vascularized, 2) a primary target organ of insulin-mediated glucose uptake that sensitively manifests insulin resistance when deficits are present, and 3) experimentally accessible using the intravital microscopy technique.

The aim of this study was to elucidate the physiological role of EC itgβ1 in controlling the skeletal muscle microcirculation and its functional importance for glucose uptake during insulin stimulation. An obstacle in understanding the physiology of EC integrins is the poor health and lethality of genetic models in which the EC itgβ1 subunit is deleted (20–24). In the present study, a mouse model was developed in which EC itgβ1 was downregulated to the threshold beyond which health is compromised. To achieve this, mice with tamoxifen (TMX) inducible EC-specific deletion of itgβ1 were generated. To avoid developmental abnormalities caused by itgβ1 deletion, TMX was administered in fully developed and mature mice at 10-weeks-of age. This model enabled us to analyze whether a quantitative loss of EC itgβ1 impairs microcirculatory function thereby creating resistance to insulin-mediated glucose uptake in otherwise healthy mice.

## Methods

Procedures involving mice were approved and carried out in compliance with the Vanderbilt University Institutional Animal Care and Use Committee. Vanderbilt University is accredited by the Association for Assessment and Accreditation of Laboratory Animal Care International. Mice were kept on a 12 hours light cycle and the environmental temperature at which mice were housed and experiments were performed was between 21-23°C.

### Animals

All mice were on the C57BL/6J strain and fed standard chow diet (5001 Laboratory Rodent; LabDiet). Mice homozygous for the lox-p flanked gene *itgβ1* (itgβ1^fl/fl^) (33) were crossed with the stem cell leukemia (SCL) promoter driven tamoxifen-inducible Cre recombinase-estrogen receptor fusion protein (CreER) transgene (34, 35). This construct causes EC-specific gene deletion (36, 37) upon tamoxifen administration. Itgβ1^fl/fl^ were used as controls. At 10 weeks old, itgβ1^fl/fl^ and itgβ1^fl/fl^SCLcre male mice were administered TMX (Sigma-T5648) reconstituted in corn oil vehicle (Sigma-C8267) at 2 mg/Kg body weight once per day for 5 consecutive days to generate mice lacking itgβ1 in EC (itgβ1^fl/fl^SCLcreTMX). One cohort of animals underwent exercise testing beginning 10 days after the last TMX dose (Fig. 1). A second cohort of animals had venous catheters implanted in a jugular vein three days post-TMX and one week prior to intravital microscopy experiments as described previously (38) for introduction of fluorescent probes during the experiment. Prior catheter implantation has the advantage of eliminating surgical stress and potential movement of the mouse on the imaging platform. A third cohort of animals had arterial and jugular vein catheters implanted at 3 days post TMX dose and underwent hyperinsulinemic-euglycemic (insulin) clamps as described previously (39) at 10 days post-TMX to determine whether a decrease in EC itgβ1 impedes whole-body insulin action. A fourth group of mice underwent terminal measurements of pulmonary function or cardiac function 10 days post-TMX, as described below.

**Figure 1.**
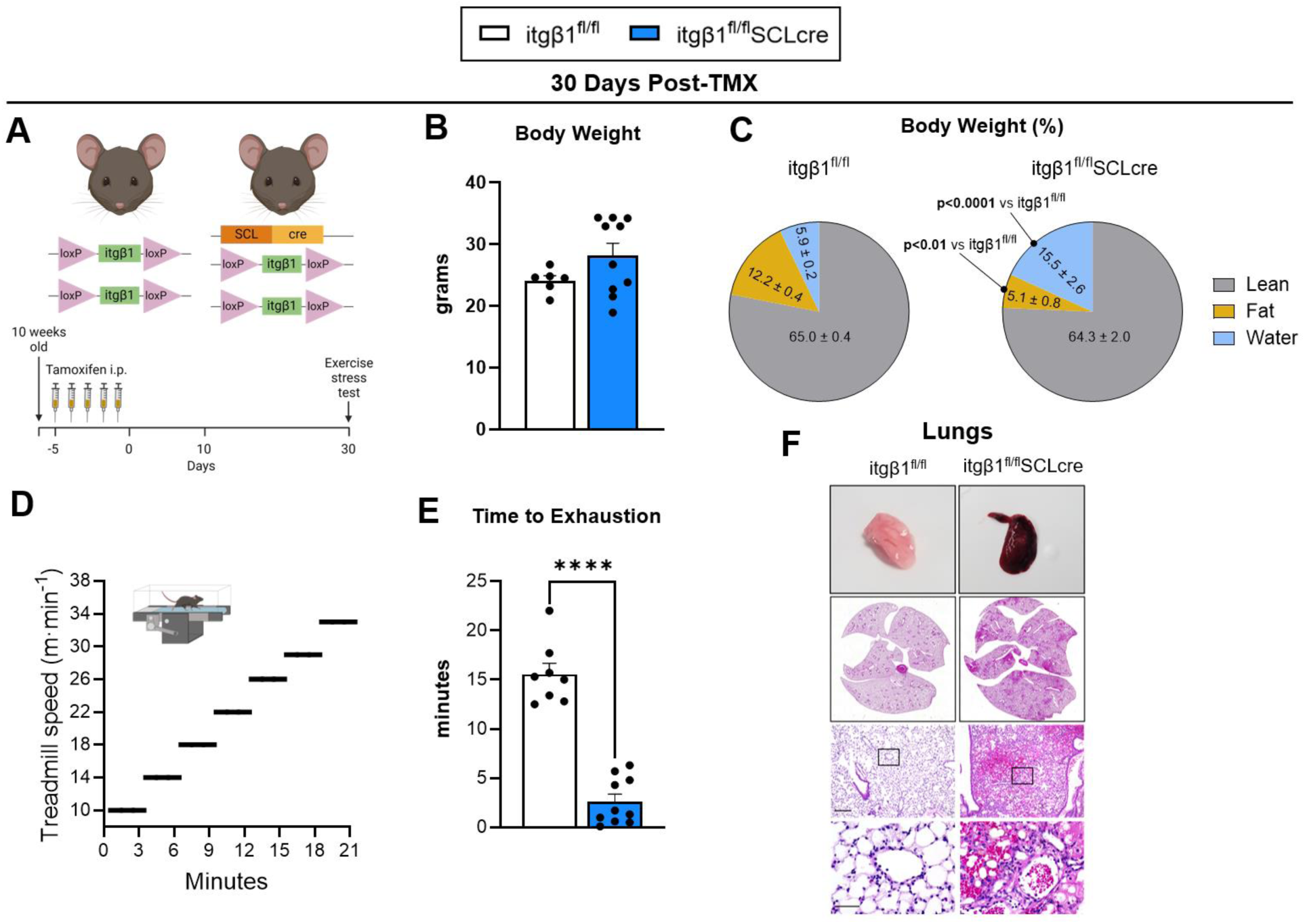
– Deletion of EC itgβ1 in adult mice is lethal. **A.** Schematic of the treatment regime used to downregulate Itgβ1 in endothelial cells (EC). itgβ1^fl/fl^ mice were crossed with mice expressing the stem cell leukemia (SCL) promoter driven tamoxifen-inducible CreER transgene. At 10 weeks old, itgβ1^fl/fl^ and itgβ1^fl/fl^SCLcre male mice were administered tamoxifen (TMX) for 5 consecutive days. **B, C**. Body weight and body composition as a percent of body weight was measured at 30 days post-TMX treatment. Data in panel C are presented as pie graphs with means and standard error for lean, fat, and water % included within each respective slice. **D**. Incremental exercise test procedure. **E**. Mice were subjected to an exercise stress test until they were no longer able to match the running speed of the treadmill for >5 seconds. This was defined as the time to exhaustion. **F**. Gross image of a single left lung lobe indicating hemorrhage in an itgβ1^fl/fl^SCLcre mouse. H&E micrographs of representative lung sections 30 days post-TMX showing widespread pulmonary hemorrhage in itgβ1^fl/fl^SCLcre mice. Cartoons in panels A and D were generated using Bioredner.com. Values are mean ± SE. n=3-8 mice/group. Panel C: Two tail student t test were used to determined statistical significance. ***p<0.001, ****p<0.0001.

### Incremental Exercise Stress Test

Stress tests were conducted on a single-lane treadmill from Columbus Instruments beginning at a speed of 10 m·min^-1^. Speed was increased by 4 m·min^-1^ every three minutes until exhaustion (**Fig. 1A**). Stress tests were performed at 10 days and 30 days post-TMX.

### Body Composition

Mouse body fat, lean mass, and body water were measured by a nuclear magnetic resonance whole-body composition analyzer (Bruker Minispec). Data are expressed as percent of total body mass.

### Lung Function Measurements

Lung function measurements were quantified using the FlexiVent apparatus (SCIREQ). Mice were anesthetized with pentobarbital sodium (85 mg/kg), and an 18-gauge tracheostomy tube was secured in the trachea. Mice were mechanically ventilated using the SCIREQ FlexiVent apparatus at 150 breaths/min and a tidal volume of 10 mL/kg body weight prior to lung function measurements. Respiratory system resistance, elastance and compliance were captured using the FlexiVent “Snapshot model”.

### Echocardiography

Transthoracic echocardiography was performed using a system (Sonos 5500, Agilent, Andover, MA) with a 15-MHz high frequency linear transducer at a frame rate of 100 frames/sec. All images were acquired at a depth setting of 20 mm. The mouse was picked up at the nape of its neck, and an ultrasound-coupling gel was applied to the precordium with the ultrasound probe. Two-dimensional targeted M-mode echocardiographic images were obtained at the level of the papillary muscles from the parasternal short-axis views and recorded at a speed of 150cm/s (maximal temporal resolution) for measurements of heart rate. All other measurements were made on-screen using the digitally recorded signals.

### Histology and Microscopy

Gastrocnemius was excised 10 days post-TMX and fixed in 4% paraformaldehyde for one hour, then in 30% sucrose overnight at 4°C. After overnight fixation, tissues were sectioned. Immunofluorescent staining for CD31 and itgβ1 was performed with anti-Integrin-β1 rabbit mAB (Cell Signaling #34971) and anti-CD31 mouse (Abcam ab24590). Secondary antibodies were goat anti-rabbit AlexaFluor 568 (ThermoFisher A-11011) and goat anti-mouse AlexaFluor 488 (ThermoFisher A-10680). A second cohort of mice was sacrificed at 28 days post-TMX. Whole mouse lungs were fixed overnight with 10% formalin prior to paraffin-embedding and staining sections with H&E. H&E sections were then imaged using a brightfield microscope. Whole muscle imaging was also performed. itgβ1^fl/fl^TMX and itgβ1^fl/fl^SCLcreTMX mice were perfused and fixed with lectin-LEA (#L32472, Invitrogen) and anti-itgβ1 rabbit mAB (#MA1-06906, Invitrogen) and 4% paraformaldehyde. SHIELD-based whole-muscle tissue clearing (40) and labeling was used to visualize vasculature and itgβ1 labeling in the gastrocnemius muscle of itgβ1^fl/fl^TMX and itgβ1^fl/fl^SCLcreTMX mice. Excised gastrocnemius muscles were fixed for an additional 24 h at 4°C, transferred to SHIELD-OFF solution (LifeCanvas Technologies, Cambridge, MA), and incubated at 4°C for 3 days. Muscles were next incubated in SHIELD-ON solution (LifeCanvas Technologies) for 24 h at 37°C. SHIELD-fixed muscles were passively cleared for 7 days at 42°C using Clear+ delipidation buffer (LifeCanvas Technologies), followed by batch labeling in SmartBatch+ (LifeCanvas Technologies) with 200µl lectin-LEA (#L32472, Invitrogen) and 6µl anti-itgβ1 rabbit mAB (#MA1-06906, Invitrogen) per muscle. Fluorescently conjugated secondary antibodies were applied in 1:2 primary/secondary molar ratios (Life Technologies, Carlsbad, CA). Labeled samples were incubated in EasyIndex (LifeCanvas Technologies) for refractive index matching (n=1.52) and imaged with SmartSPIM (LifeCanvas Technologies) at 2µm z-step and 1.8µm pixel size (24, 25). Following acquisition, images were stitched to generate composite TIFF images using a modified version of Terastitcher (Bria et al., 2019). Stitched TIFF images were converted to Imaris files using Imaris File Converter 9.2.1 and 3D renderings of muscles were visualized using Imaris software (version 9.5.1; Bitplane, Zurich, Switzerland).

### Intravital Microscopy

Intravital microscopy was performed in mice as previously described (5, 6, 41). Studies were performed under isoflurane anesthesia (SomnoSuite; Kent Scientific). Doses of 2 and 1.5% isoflurane were used for induction and maintenance of anesthesia, respectively. The lateral gastrocnemius was exposed for visualization by trimming away skin and fascia and then placed on a glass coverslip immersed in 0.9% saline. Body temperature of 37°C was maintained using a homeothermic heating blanket and temperature probe (Harvard Apparatus). Approximately 8 mg/kg (100 μL injection volume) of a tetramethylrhodamine (TMR)-labeled 2-megadalton (MDa) dextran (Thermo Fisher) was infused through the jugular vein catheter. A 2 MDa molecule remains in the lumen of capillaries with high resistance endothelial walls and used to identify microcirculatory structure. Imaging began at t=0 min and continued until 10 minutes after the TMR bolus and insulin bolus, respectively. The focal plane for imaging was maintained using the Perfect Focus System on the Nikon Eclipse Ti-E (Nikon Instruments). Plasma fluorescence was excited using a Sola Light Engine LED lamp (Lumencore) and visualized with a Plan Apo 10× objective (Nikon Instruments). Images were recorded at 100 fps using a 10 ms exposure time with a Flash 4.0 sCMOS camera (Hamamatsu). High frame rate imaging was achieved by directly streaming time-lapse experiments to computer RAM using a Camlink interface (Olympus). Videos underwent real-time 4 × 4-pixel binning during acquisition to improve signal/noise ratio, and image brightness was manually adjusted post hoc in NIS Elements (Nikon Instruments) to ensure an average pixel intensity of approximately 0.5 with a minimal extent of over/underexposed regions prior to export in a MATLAB-readable format. Capillary flow velocity, hematocrit, and density were quantified as previously described (38). Briefly, 5 s videos of the gastrocnemius microcirculation acquired at 100 frames·s^-1^ were processed to remove motion artifacts, identify in-focus capillaries, and track the motion of red blood cells (RBCs) based on the shadows they produce in plasma fluorescence. Perfusion metrics included mean capillary flow velocity (MFV; µm·s^-1^), proportion of perfused vessels (PPV; fraction of total capillary density with any detectable flow during the 5-s video), and perfusion heterogeneity index (PHI; a unitless measure of spatial flow variability). PHI was defined as the natural log of *V̅*_max_ -*V̅*_min_ divided by the MFV. *V̅*_max_ and *V̅*_min_ represent velocity in the fastest and slowest flowing capillaries, respectively. Five fields of view were captured for each mouse, and MFV and PHI represent an average of all five acquisitions.

### Transendothelial Insulin Efflux

For imaging of transendothelial insulin efflux (INS-647), rho-dex fluorescence was excited using light from a 561-nm solid-state laser and detected on a multichannel PMT. The near-infrared fluorophores were excited using a helium-neon 633-nm laser, and emitted light was detected using a GaAsP detector. For both rho-dex and the near-infrared fluorophores, excitation and emission light were passed through an MBS 488/561/633 dichroic mirror. A confocal pinhole was set to give an optical section of 8 μm (±4 μm about the focal plane). Imaging of the rho-dex and near-infrared fluorophores was performed using 2-channel sequential excitation and detection to prevent bleed-through. Switching between channels occurred every line to minimize channel mis-registration due to intra scan motion artifacts. Eight-bit-intensity, 1,024-by-1,024-pixel images were acquired with unidirectional scanning. For each time point, a 4-slice Z-stack was acquired in each channel using a step size of 4 μm to avoid aliasing. PMT settings were adjusted to maximize the dynamic range of the image and kept constant for each given experiment to allow for quantitative comparisons. After the selection of an imaging region but before administration of the probes, a background image was acquired. Subsequently, INS-647 (2 U/kg body weight) was infused through the indwelling vein catheter and followed with a 20-μl pulse of saline. Images were then acquired using the procedure described above every minute for the first ten minutes after probe injection.

Intravascular and extravascular spaces were segmented in raw images using a custom ImageJ (NIH) macro. A mask of vascular structures in the rho-dex channel was created using Otsu thresholding. This mask was then closed (dilation then erosion) and opened (erosion then dilation) with mathematical morphometric techniques. We then applied a median filter (radius = 4 pixels) to remove any remaining noise. Object recognition was then performed with the ImageJ Particle Analyzer to select vessel structures that were larger than 50 μm^2^ in area and had circularity from 0 to 0.5. The final mask of these vascular structures was then applied to the INS-647 channel to measure the intravascular intensity of INS-647. The mask was then dilated in steps of 0.5 μm out to 3 μm from the capillary wall to segment the interstitial spaces immediately adjacent to the capillary. The intensity of INS-647 was then measured in all of these extravascular segments. The contribution of INS-647 from multiple vessels to a single interstitial segment was avoided by (a) imaging of regions in which capillaries were not immediately adjacent to one another (i.e., >15 μm apart in most cases) and (b) restriction of analysis of the interstitial space to 3 μm from the capillary. Following the extraction of intravascular and extravascular INS-647 intensity versus time profiles, a number of postprocessing steps were performed. First, data were background-subtracted using intensity values from background images collected during imaging experiments. Subsequently, the loss of fluorescence due to photobleaching was corrected for each scan in the interstitial spaces. We chose not to correct the intravascular space for photobleaching because intravascular INS-647 is circulating through the capillary blood stream and therefore most likely does not reside in the field of view long enough to undergo significant bleaching. The intensities from the 4 slices of the Z-stack were averaged to give a single value at each point in time and space.

### Hyperinsulinemic-euglycemic Clamp Procedure

Catheters were surgically implanted in a carotid artery and jugular vein for sampling and infusions, respectively, one week before clamp experiments (39). Mice were transferred to a 1.5 L plastic container without access to food 5 hour prior to the start of an experiment. Glucose clamps were conducted as described previously (39). Mice were neither restrained nor handled during clamp experiments. [3-^3^H]glucose was primed and continuously infused from t=-90 min to t=0 min (0.06 µCi·min^-1^). The insulin clamp was initiated at t=0 min with a continuous insulin infusion (4 mU·kg^-1^·min^-1^) and variable glucose infusion initiated and maintained until t=155 min. The glucose infusate contained [3-^3^H]glucose (0.06 µCi·µl^-1^) to minimize changes in plasma [3-^3^H]glucose specific activity. Arterial glucose was monitored every 10 min to provide feedback to adjust the glucose infusion rate (GIR) as needed to maintain euglycemia. Erythrocytes were infused at a rate calculated to compensate for blood withdrawal over the duration of the experiment. [3-^3^H]glucose specific activity was determined at −15 min and −5 min for the basal period, and every 10 min between 80 to 120 min for the clamp period to assess glucose disappearance (Rd) and endogenous glucose production (EndoRa). A 13 µCi intravenous bolus of 2-[^14^C]-deoxyglucose ([^14^C]2DG) was administered at 120 min to determine the glucose metabolic index (Rg), an index of tissue-specific glucose uptake (42–44). A modified Rg (Rgʹ) was calculated by gaining a measure of individual tissue decay curves by multiplying the arterial decay curve by the tissue to arterial [^14^C]2DG ratio (the ratio is unitless) measured in the terminal sample (t=145 min), as previously described (45). Blood samples were collected at 122, 125, 130, 135 and 145 min to measure plasma [^14^C]2DG. At 145 min, mice were euthanized, and tissues immediately harvested and freeze-clamped to measure the tissue accumulation of 2-[^14^C]-deoxyglucose-phosphate ([^14^C]2DGP). Detailed methodology is available via VMMPC webpage https://vmmpc.org.

### Skeletal Muscle Insulin Signaling

Immunoblotting was performed on vastus lateralis muscle post insulin clamp. Immediately following the clamp procedure, skeletal muscle was excised and snap frozen for subsequent analysis. Mice that had not been stimulated with insulin were used as negative controls. Triton X-100 tissue lysates (5-15 µg/lane) were analyzed by western blot for total or activated levels of the following proteins: rabbit anti-AKT, #9272; rabbit anti-phospho AKT_Ser473_, #4060; rabbit anti-ERK, #9102; rabbit anti-phospho ERK_Thr202/Tyr204_, #9101; rabbit anti-GSK3β, #9315; rabbit anti-phospho GSK3β_Ser9_. #5558; Cell Signaling). After incubation with appropriate HRP-conjugated secondary antibodies (rabbit anti-IgG, #7074, Cell Signaling) bands were detected via chemiluminescence. Intensity of individual bands were quantified using Image Lab^TM^ (version 6.0.0, Bio-Rad Laboratories, Inc.), and expressed as a ratio to total protein stain per membrane using amido black staining. For phosphorylated protein quantification, band intensities were expressed as a ratio to the total protein (e.g. pAKT/AKT). Values are expressed as fold-difference with the vehicle group set to 1.

### Ex Vivo 2-Deoxy-D-Glucose Uptake

Isolated soleus muscle 2-deoxyglucose uptake was measured as previously described (46). After a 15-min basal incubation period, muscles were transferred to fresh media and incubated for 20 min in the absence or presence of insulin (100 nmol/l). After the 20 min stimulation, 2DG uptake was measured for 10 min in fresh media containing 2-deoxy-d-glucose (1 mmol/L) and 2-[1-^14^C]deoxy-d-glucose (0.25 μCi/mL). Muscles were then lysed, and 2-[1-^14^C]deoxy-d-glucose-phosphate radioactivity was measured.

### Statistical Analysis

Student’s t-tests were run for between group comparisons. If data did not follow a Gaussian distribution, non-parametric Mann-Whitney tests were used to determine statistical significance. In experiments that contained more than two groups, one-way analysis of variance (ANOVA) or two-way ANOVA models were used with pairwise comparisons using Tukey or Sidak correction. Brown-Forsythe correction was applied to groups with unequal variance. Data are presented as mean ± standard error (SE). A p value of <0.05 was used to determine significance.

## Results

### Deletion of EC itgβ1 in adult mice is lethal

Inducible deletion of the EC itgβ1 subunit between postnatal day 2 and day 30 causes premature death, characterized by vascular instability and hemorrhage (20). We hypothesized that deletion of the EC itgβ1 subunit in fully developed adult mice may also be lethal, however, the point of unrecoverable health was unknown. At 10 weeks old, itgβ1^fl/fl^ and itgβ1^fl/fl^SCLcre mice were administered TMX for 5 consecutive days (**Fig. 1A**) to generate EC-specific deletion of the itgβ1 subunit. We measured body mass and body composition at 4 weeks (28-30 days) post-TMX. Body mass and the percentage of lean mass were not different between itgβ1^fl/fl^SCLcreTMX and itgβ1^fl/fl^TMX mice (**Fig. 1B&C**). The percentage of fat mass was decreased in itgβ1^fl/fl^SCLcreTMX mice, whereas percent water weight was nearly doubled at ∼30 days post-TMX with the presence of edema (**Fig. 1C**). The ability to engage in incremental exercise was implemented as an indicator of health because exercise requires high functioning cardiovascular, respiratory, and endocrine systems to meet the metabolic demands of sustained physical exertion. At 28-30 days post TMX, itgβ1^fl/fl^SCLcreTMX and itgβ1^fl/fl^TMX mice were subjected to incremental exercise stress tests (**Fig. 1D**). Severely impaired exercise tolerance manifested in itgβ1^fl/fl^SCLcreTMX mice (**Fig. 1E**). At this time point (≈30 days post-TMX), itgβ1^fl/fl^SCLcreTMX mice were moribund. In compliance with ethical treatment of animals, mice were immediately euthanized if body weight loss >15% in a week, lack of alertness, and respiratory distress were noted. Necropsy of lung tissue revealed pulmonary hemorrhage due to damage to the pulmonary capillaries in the itgβ1^fl/fl^SCLcreTMX mice (**Fig. 1F**). This pathology was the diagnosed cause of death.

We then set out to define the time point post-TMX when itgβ1 in skeletal blood vessels was deleted but there was no decline in health. This was found to be 10 days after the last dose of TMX at which time there was a ∼30% knockdown of itgβ1 in CD31 positive ECs in itgβ1^fl/fl^SCLcreTMX mice (**Fig. 2A-D**). This was confirmed by immunofluorescence by microscopy of skeletal muscle tissue and in intact whole vessels using light sheet microscopy by visualizing the intensity of co-localization of itgβ1 and individual capillaries (**Fig. 2E**). The reduction in CD31 signal intensity was quantified relative to area and is indicative of reduced capillary density (**Fig. 2A&C**). At this time point, we found that itgβ1^fl/fl^SCLcreTMX mice have equivalent pulmonary function and cardiac function compared to itgβ1^fl/fl^ mice (**Fig. 3A-C**). Itgβ1^fl/fl^SCLcreTMX mice showed similar body weight and percent lean mass at 10 days post-TMX (**Fig. 4A-B**). There was a small reduction in the percent fat mass, and a small increase in the percentage of body water in itgβ1^fl/fl^SCLcreTMX mice at 10 days post-TMX (**Fig. 4B**). Consistent with pulmonary and cardiac functions, exercise tolerance was comparable between itgβ1^fl/fl^SCLcre and itgβ1^fl/fl^ mice (**Fig. 4C**). These data show that at 10 days post-TMX, itgβ1^fl/fl^SCLcre and itgβ1^fl/fl^ mice are phenotypically similar at rest and during the metabolic demands imposed by exercise. We therefore decided to conduct studies of microcirculation and insulin action at the 10 days post-TMX time point in itgβ1^fl/fl^SCLcreTMX and itgβ1^fl/fl^TMX mice.

**Figure 2.**
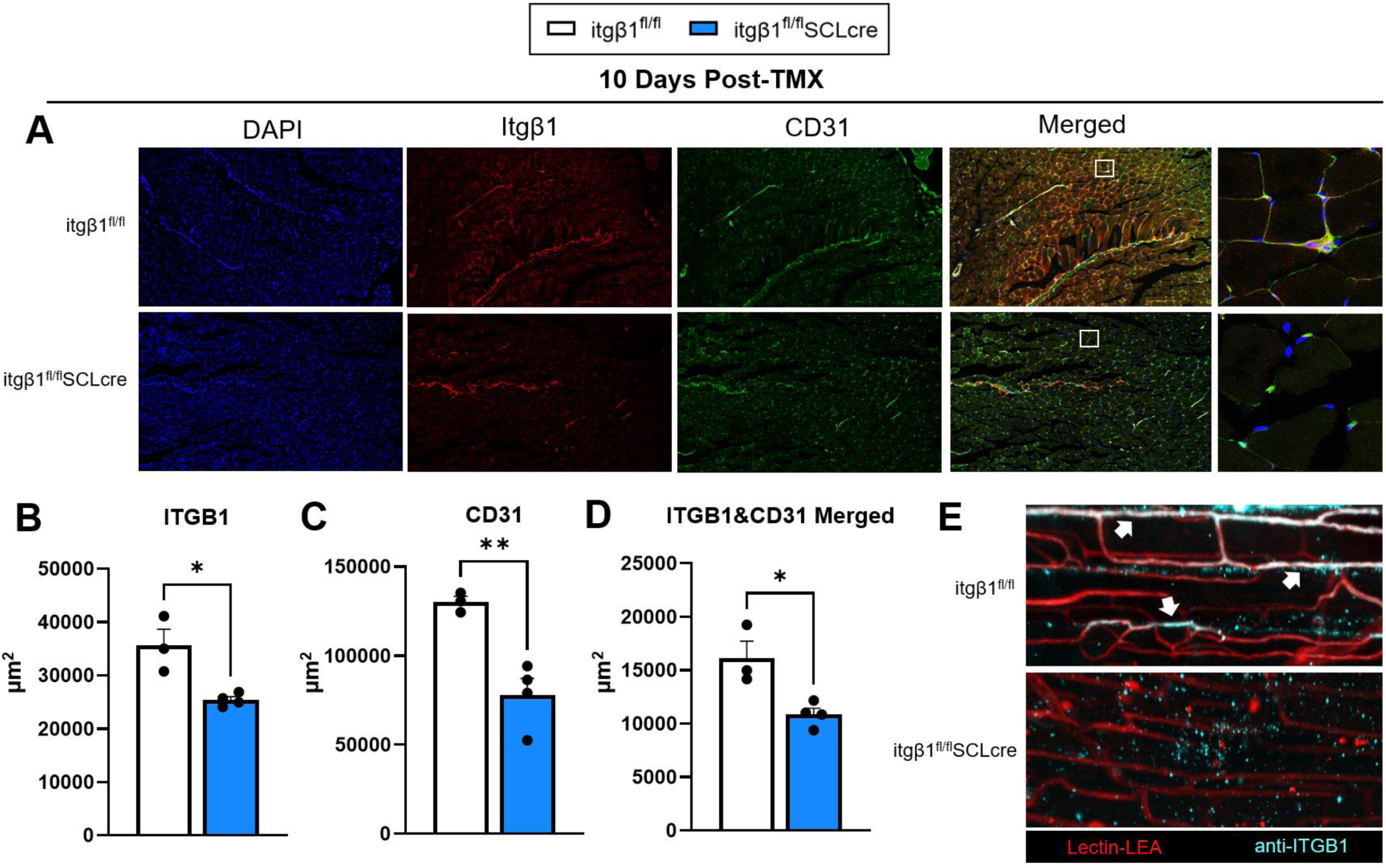
– Downregulation of EC-itgβ1 decreases capillary density in skeletal muscle. **A.** Representative immunofluorescence images of gastrocnemius muscle at 10X, with 63X merged inset (far right panel) stained with ITGB1, CD31, and DAPI. **B**. Immunofluorescence microscopy shows that itgβ1 is decreased in the skeletal muscle of itgβ1^fl/fl^SCLcre mice 10 days after the last dose of TMX. **C.** CD31 is decreased in the skeletal muscle of itgβ1^fl/fl^SCLcre mice. **D.** Co-localization of CD31 and itgβ1 shows an overall decrease in the gastrocnemius muscle of itgβ1^fl/fl^SCLcre mice. **E.** Representative images of individual capillaries (Lectin-LEA) and anti-ITGB1 co-expression in itgβ1^fl/fl^SCLcre and itgβ1^fl/fl^ mice. Arrows indicate strong ITGB1 signal colocalized with Lectin positive capillaries. Panel E representative images were taken using the Zeiss LSM710 confocal microscope. Two tail student t test were run to compare groups. Three to four mice per genotype were used to generate sections and images. The average fluorescent intensity across multiple fields of view (FOV) were conducted to generate a single mean, resulting in a biological n=3-4/group. Data are mean ± SE. *p<0.05; **p<0.01.

**Figure 3.**
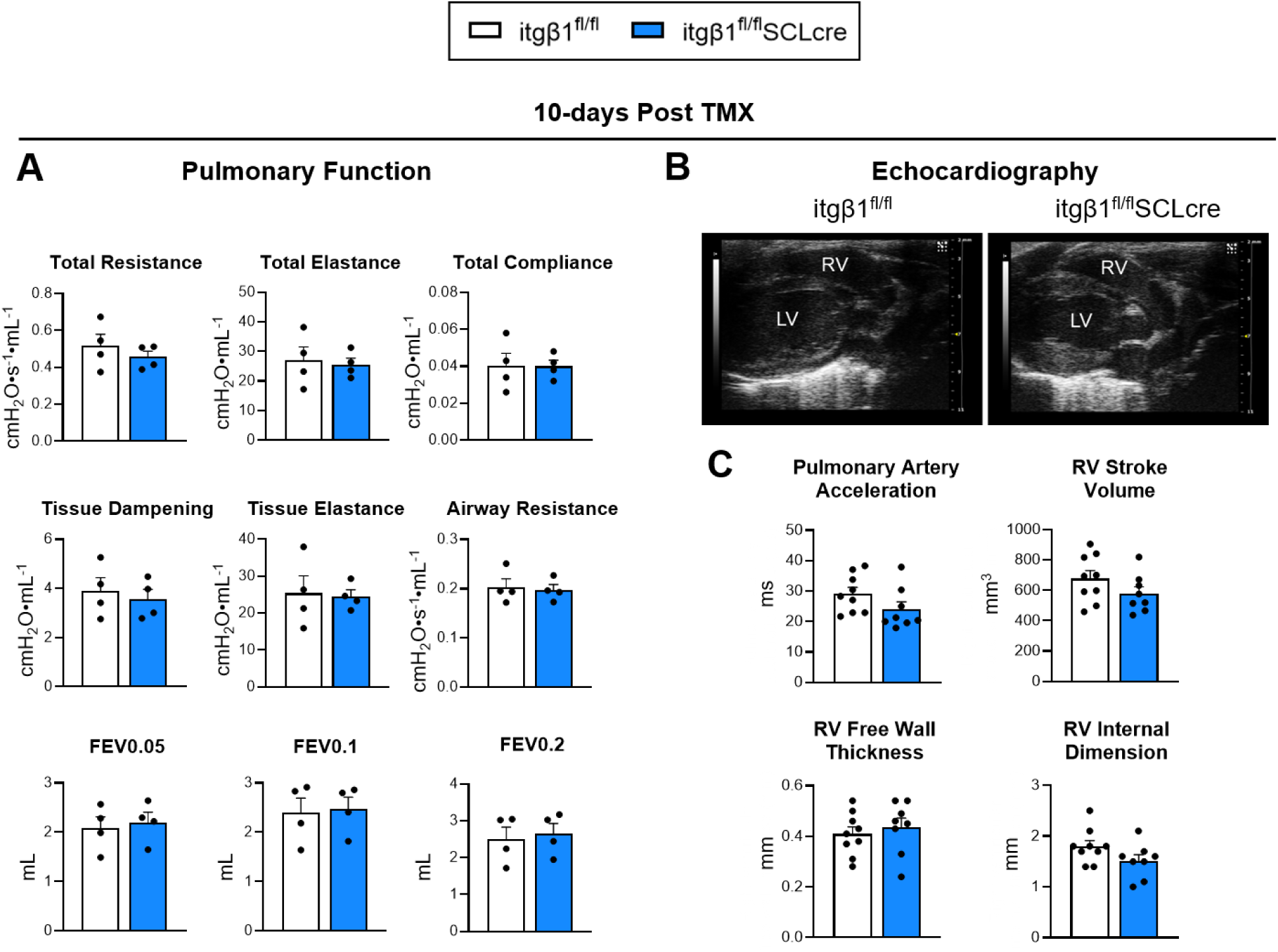
– EC itgβ1 downregulation does not affect pulmonary or cardiac function at 10 days post-TMX. **A.** In vivo respiratory gas analysis was performed 10 days post TMX. No differences in pulmonary function were found between groups. **B**. Echocardiography was performed 10-days post TMX. A representative image showing the left ventricle (LV) and right ventricle (RV) of the myocardium. Real-time video images were analyzed to quantify **C**. quantify pulmonary artery acceleration, right ventricular stroke volume, right ventricular wall thickness, and right ventricular internal dimension. These parameters were similar between genotypes. Rrs, resistance of the total respiratory system; Ers, elastance of the total respiratory system; Crs, compliance of the total respiratory system; Rn, airway resistance; G, tissue dampening; H, tissue elastance; FEV0.5, forced expiratory volume in 0.05 sec; FEV0.1, forced expiratory volume in 0.1 sec; FEV0.2, forced expiratory volume in 0.2 sec. 10x image scale bar = 200 μm. 60x image scale bar = 50 μm. Two tail student t test was used to compare groups. Data are mean ± SE. n=4-9 mice/group.

**Figure 4.**
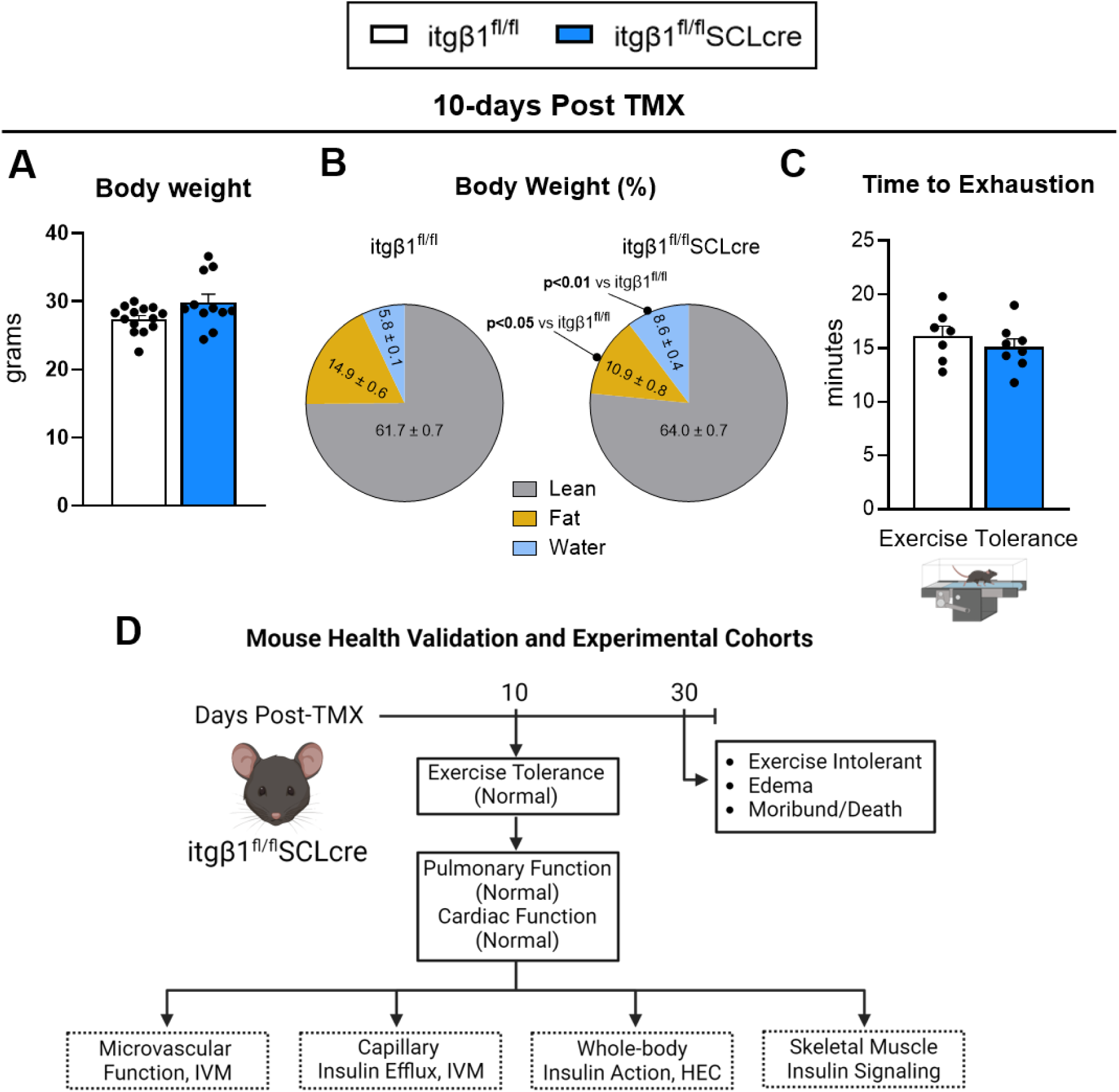
– Downregulation EC itgβ1 does not affect body weight or exercise tolerance at 10 days post-TMX. At 10 weeks-of-age itgβ1^fl/fl^SCLcre and itgβ1^fl/fl^ mice were administered tamoxifen (TMX) for 5 days. Ten days after the last TMX treatment, **A**. body weight and **B**. body composition were measured **C**. Mice were subjected to an exercise stress test to evaluate whole-body physical function. The treadmill speed increased every 3 minutes until mice could no longer maintain running speed, indicating exercise exhaustion. **D**. Summary of health validation between 10 days and 30 days post-TMX. At 10 days post-TMX itgβ1^fl/fl^SCLcre mice have normal exercise tolerance, cardiac and pulmonary function. By 30 days post-TMX, these mice are severely exercise intolerant and moribund. Panel D was generated via Biorender.com Two tail student t test were run to compare groups. n=7-13 mice/group. Data are mean ± SE.

### EC itgβ1 regulates microvascular hemodynamics without affecting capillary insulin efflux

EC integrins are postulated to determine microcirculatory blood flow, capillary surface area, and structural integrity of capillaries (27, 47, 48). These characteristics determine the delivery of both perfusion-limited molecules (e.g. glucose, O_2_) and dispersion-limited molecules (e.g. insulin). We tested the hypothesis that skeletal muscle EC itgβ1 is necessary for microcirculatory function. Microvascular hemodynamics were quantified using intravital microscopy (see Methods). Thirty percent loss of EC itgβ1 reduced mean capillary flow velocity by ∼30% (**Fig. 5A**). The reduced capillary blood flow corresponded to reduced proportion of perfused vessels (**Fig. 5B**) and greater flow heterogeneity and between capillaries (**Fig. 5C)** in itgβ1^fl/fl^SCLcreTMX compared to itgβ1^fl/fl^TMX mice. Hematocrit variability reflects the movement of particulates in a capillary bed. There was more hematocrit variability (p=0.051) in itgβ1^fl/fl^SCLcreTMX mice (**Fig. 5D**). We then tested the hypothesis that knockdown of EC itgβ1 alters the capillary insulin efflux by monitoring the capillary and peri-capillary interstitial fluorescence insulin (INS-647) using IVM. The capillary insulin efflux was similar between genotypes as neither the gradient decay constants nor the fractional capillary removal were significantly different (**Fig. 5E&F**). These data show that a 30% reduction in EC itgβ1 reduces the rate of capillary perfusion but does not significantly disrupt transcapillary insulin efflux in skeletal muscle.

**Figure 5.**
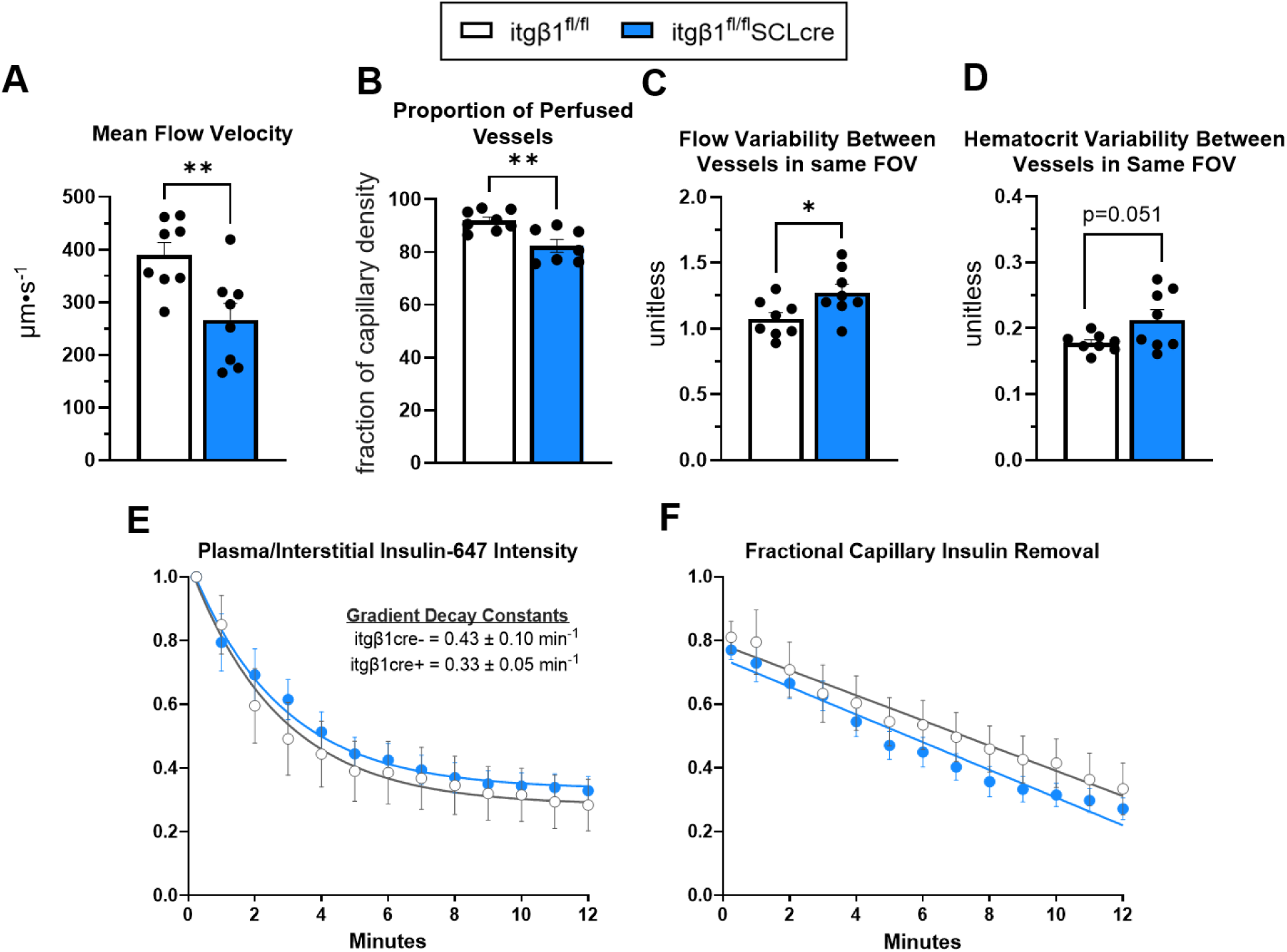
– EC itgβ1 regulates microvascular hemodynamics without affecting capillary insulin efflux. Intravital microscopy (IVM) was performed in anesthetized mice 10 days post-TMX. **A.** A decrease in flow velocity through skeletal muscle capillary beds in itgβ1^fl/fl^SCLcre mice was observed. **B.** The proportion of perfused vessels was reduced in itgβ1^fl/fl^SCLcre mice than itgβ1^fl/fl^ mice. **C.** Within one field of view (FOV), there was significantly more heterogeneity in flow velocity when comparing all vessels in the itgβ1^fl/fl^SCLcre mice than the itgβ1^fl/fl^ mice. **D.** Within one FOV, there was more heterogeneity in hematocrit distribution in the itgβ1^fl/fl^SCLcre than itgβ1^fl/fl^ mice. **E**. Insulin efflux was determined via intravital microscopy following an INS-647 bolus. T = 0.25 minutes indicates the beginning of imaging, which occurs ∼15 seconds after injection of INS-647. Interstitial space is defined as the region emanating 1-3 µm from the capillary wall. Ratio of plasma to interstitial INS-647 as a function of time following INS-647 injection, normalized to the ratio at t = 0 minutes. Decay constant of the plasma/interstitial INS-647 ratio, a measure of transendothelial insulin transport kinetics. **F**. The fractional removal rate of capillary insulin. Two tail student t test was used to compare groups. n=6-9/group. Data are mean ± SE. *p<0.05; **p<0.01.

### Glucose uptake during a hyperinsulinemic-euglycemic insulin clamp is dependent on EC itgβ1

Skeletal muscle capillary flow rate and the area for capillary exchange are increased during the metabolic demands of insulin (6). This is important for both insulin and glucose delivery to target muscle. We tested whether decreased microvascular perfusion that manifests in itgβ1^fl/fl^SCLcreTMX mice corresponded to deficits in glucose kinetics during the metabolic demands of an insulin clamp (**Fig. 6A**). Steady state glucose infusion rate was suppressed by ∼25% in itgβ1^fl/fl^SCLcreTMX mice indicating a reduction in whole-body insulin action (**Fig. 6B**). Fasting endogenous glucose appearance (EndoRa) and the rate of glucose disappearance (Rd) were lower in itgβ1^fl/fl^SCLcreTMX compared to itgβ1^fl/fl^TMX mice (**Fig. 6C**). itgβ1^fl/fl^SCLcreTMX and itgβ1^fl/fl^TMX mice both exhibit complete suppression of glucose production during the insulin clamp (**Fig. 6C**). Glucose disappearance was decreased in itgβ1^fl/fl^SCLcreTMX mice during the insulin clamp, suggesting non-hepatic reduction in insulin action (**Fig. 6D**). Fasting insulin was not different and increased similarly in itgβ1^fl/fl^SCLcreTMX and itgβ1^fl/fl^TMX mice during the procedure (**Fig. 6E**). A bolus of [^14^C]2DG was administered during the clamp to determine the glucose metabolic index (Rgʹ), an index of tissue-specific glucose uptake. Skeletal muscle, heart, and adipose tissue Rgʹ, were reduced in itgβ1^fl/fl^SCLcreTMX mice (**Fig. 6F**). In addition, the ratio of steady state tissue water [3-^3^H]glucose to arterial plasma [3-^3^H]glucose during the insulin clamp was calculated as a measure of tissue glucose uptake. This ratio was decreased in skeletal muscle and heart, but not epididymal white adipose tissue (eWAT) or inguinal white adipose tissue (iWAT) in itgβ1^fl/fl^SCLcreTMX compared to itgβ1^fl/fl^TMX mice (**Fig. 6G**). These data show that a reduction in EC itgβ1 impairs non-hepatic whole-body insulin sensitivity and glucose uptake.

**Figure 6.**
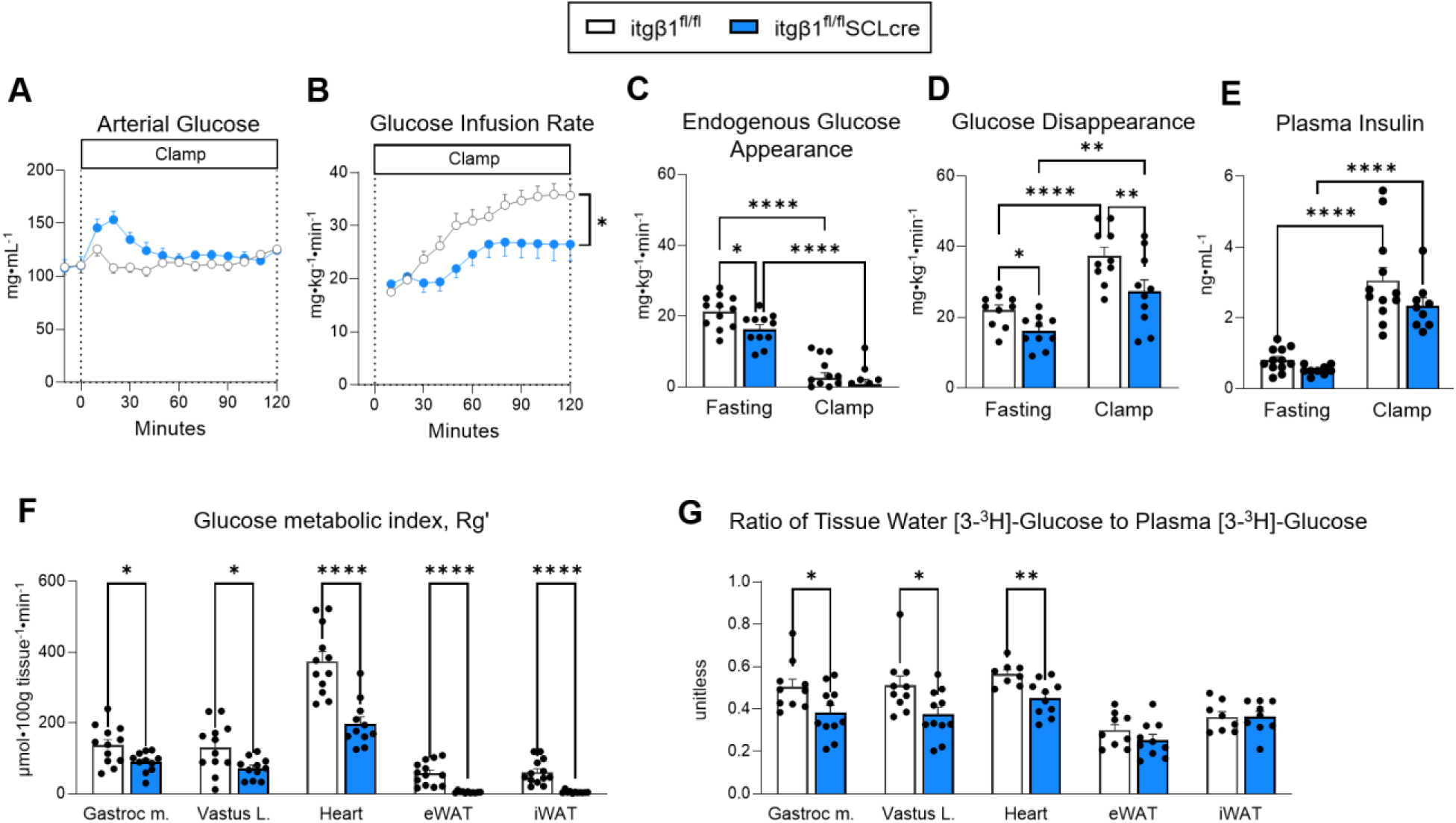
– Downregulation EC itgβ1 impairs whole-body insulin action. Hyperinsulinemic-euglycemic clamps were performed 10 days after the last TMX dose in conscious unstress mice. **A**. Arterial glucose was administered and monitored during the glucose infusion and insulin clamp. **B.** Glucose infusion rate was monitored every 10 minutes for each group. **C.** The rate of endogenous glucose appearance was measured for each group. **D**. Glucose flux was measured between groups over 100 minutes. **E**. Insulin was measured at baseline and during the clamp period. **F.** The glucose metabolic index (Rgʹ) was measured in gastrocnemius muscle, vastus lateralis muscle, epididymal (eWAT), and inguinal white adipose tissue (iWAT). **G**. Ratio of tissue water [^3^H]-glucose over plasma [^3^H]-glucose at the end of the clamp period. Panel A&B: Two-way ANOVA with repeated measures were run with Tukey adjustment. Panel C-E: Two-way ANOVA with group and condition as factors were run with Tukey adjustment. Panel F&G: Two tail student t test were run to compare groups. Data are mean ± SE n=9-12/group. *p<0.05; **p<0.01.

### Myocellular insulin signaling and ex vivo glucose uptake is not dependent on EC itgβ1

Given the decrease in whole-body insulin action, we tested whether insulin sensitivity was impaired in isolated skeletal muscle in the absence of the microcirculation. We analyzed the activation of insulin-regulated downstream kinases, including Akt, GSK3β and ERK in vastus lateralis muscle in the basal state and at the end of the insulin clamp. We observed decreased Akt_Ser473_ activation itgβ1^fl/fl^SCLcreTMX mice compared to control mice in the absence of insulin stimulation (**Fig. 7A&B**). However, at the end of the clamp, we detected increased activation of Akt, GSK3β, and ERK in both itgβ1^fl/fl^SCLcreTMX and itgβ1^fl/fl^TMX mice (**Fig. 7A-D**). Muscle insulin-stimulated 2-[1-^14^C]deoxy-d-glucose (2DG) uptake was also tested in an *ex vivo* system to assess whether itgβ1^fl/fl^SCLcreTMX mice develop an inability to respond to transport or phosphorylated glucose. Soleus muscle was excised and 2DG uptake was measured in itgβ1^fl/fl^SCLcreTMX and itgβ1^fl/fl^TMX mice in the absence or presence of insulin. (**Fig. 7E**). No differences in soleus 2DG uptake were detected between itgβ1^fl/fl^SCLcreTMX and itgβ1^fl/fl^TMX mice either in the presence or absence of insulin. Collectively, these data show that downregulation of EC itgβ1 results in decreased glucose uptake that is mediated by the microvasculature, with intact muscle insulin action.

**Figure 7.**
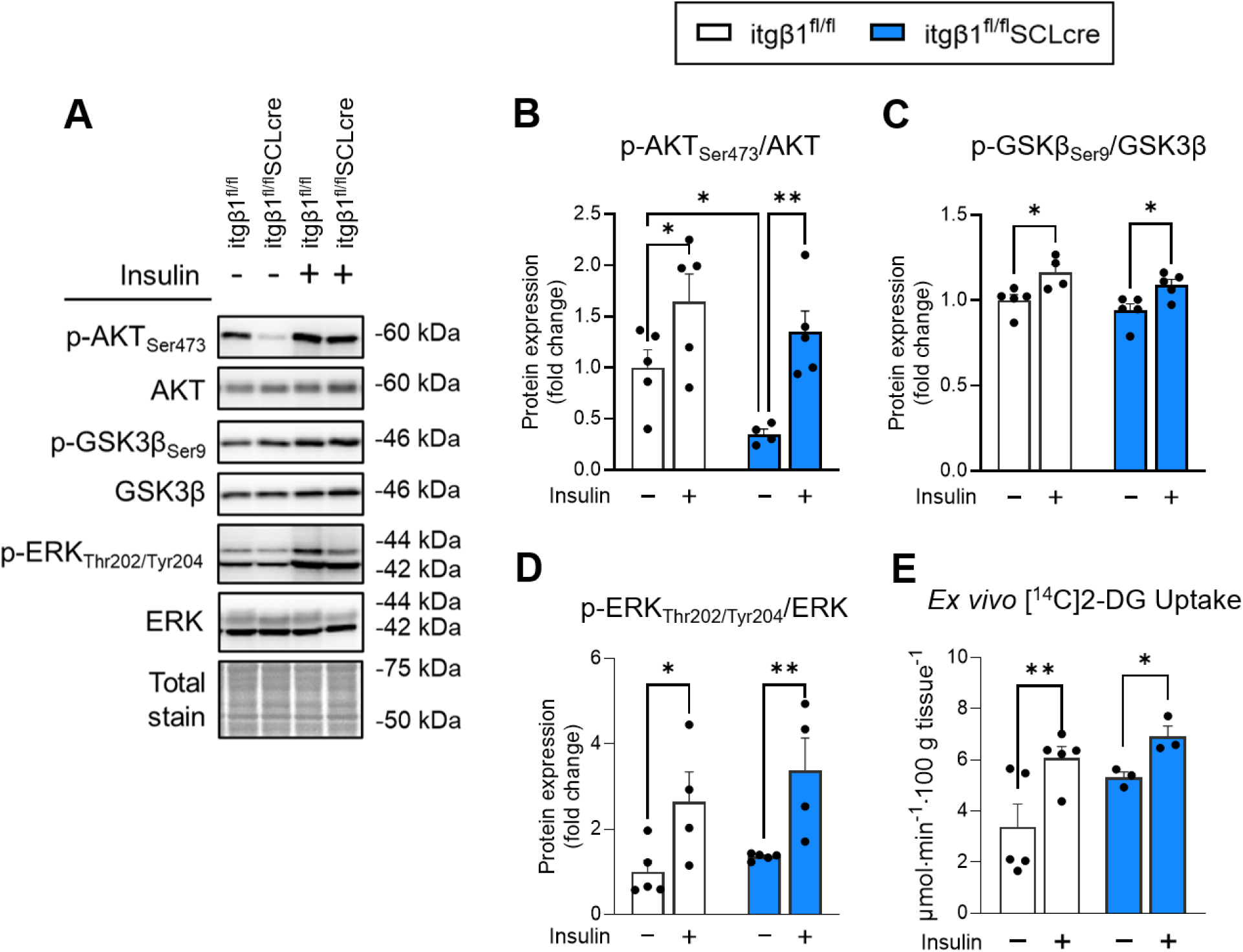
– Myocellular insulin signaling and ex vivo glucose uptake is not dependent on EC itgβ1. **A.** Lysates were prepared from gastrocnemius muscle collected from 10-day post-TMX mice and analyzed by Western blot for levels of activated and total AKT, GSK3β, and ERK under basal and insulin stimulated (clamp) conditions. **B-D.** Bands were quantified and values are expressed as activated/total proteins as fold changes relative to the basal itgβ1^fl/fl^ condition **E**. Ex vivo 2DG uptake in isolated soleus muscle in the absence or presence of insulin (100 nM). Two-way ANOVA with repeated measures were run with Tukey adjustment. Data are mean ± SE. Panels A-D n=4-5/condition. Panel E, bilateral soleus muscle was isolated from n=3-5 mice per genotype. Soleus from one leg was stimulated with insulin and the contralateral muscle with vehicle. *p<0.05; **p<0.01

## Discussion

The objective of the current study was to elucidate the physiological role of EC itgβ1 in control of the skeletal muscle microcirculation and its functional importance during insulin stimulation. This is important because the microcirculation represents a site of resistance to muscle glucose uptake that can be rate-limiting for the initiation of insulin action in muscle. To fill this knowledge gap, a tamoxifen-inducible EC itgβ1 knockdown mouse model was developed. Using robust in vivo experimental platforms (i.e., intravital microscopy and hyperinsulinemic-euglycemic clamp) we show that itgβ1^fl/fl^SCLcre mice compared to itgβ1^fl/fl^ littermates have, i) deficits in capillary flow rate, flow heterogeneity, and capillary density; ii) impaired insulin-stimulated glucose uptake despite sufficient transcapillary insulin efflux; and iii) reduced insulin-stimulated glucose uptake due to perfusion-limited glucose delivery. These data establish that EC itgβ1 is necessary for microcirculatory function and to meet the metabolic challenge of insulin stimulation. On a broader scale, our findings emphasize that endothelial integrins are potential regulatory gatekeepers of the exchange of bloodborne molecules between circulation and peripheral tissues.

Germline deletion of the primary EC itgβ subunit, itgβ1, is embryonic lethal (21, 23, 24, 49). Similarly, conditional ablation of EC itgβ1 between postnatal day 2 and day 30 also causes premature death, characterized by vascular instability and hemorrhage (20). We extend these findings to adult mice, showing that inducible deletion of EC itgβ1 in 10-week-old mice results in death within 5 weeks of TMX treatment. This was characterized by a loss of exercise tolerance and severe pulmonary edema which was diagnosed as the cause of death. The essential role of interstitial pressures as keys for survival is evident by studies that show that EC-specific deletion of the integrin α9 subunit, which binds itgβ1, in mice causes lethality due to lymphedema within 12 days of birth (50). The health of an animal is a sensitive determinant of microcirculatory function and metabolism. Therefore, the focus of our studies on microcirculatory effects on metabolism required that we determine whether there was a window in time where a detectable knockdown in the EC itgβ1 subunit could be detected in otherwise healthy mice. We found that this time point was 10 days post-TMX. Importantly, EC itgβ1 was reduced by ∼30% which was accompanied by a proportional reduction in CD31+ capillaries, a decrease in mean flow velocity, and an increase in the heterogeneity of capillary blood flow in skeletal muscle. These deficits compromise nutrient and hormone delivery to peripheral tissue (41). The loss of skeletal muscle capillaries could be attributed to reduced EC survival (23), decreased EC proliferation (25), or a combination of the two. EC apoptosis typically occurs when nutrients are limiting or survival signals such as those triggered by vascular endothelial like growth factor (VEGF) are reduced (51, 52). In the absence of itgβ1, ex vivo studies show reduced EC survival, but EC proliferation is sustained (23). Together, our findings support an indispensable role for EC integrin β1 receptors in maintaining microcirculatory function in otherwise healthy mice.

In this study, microvascular dysfunction is evident in the sedentary state in itgβ1^fl/fl^SCLcreTMX mice. The increase in cardiac output and skeletal muscle blood flow in response to exercise in EC itgβ1^fl/fl^SCLcreTMX mice overcome deficits due to reduced itgβ1 such that performance on an incremental stress test is not different than in littermate control mice at 10 days post-TMX. It is possible that energy production from nutrients stored in muscle may have been accelerated in EC itgβ1^fl/fl^SCLcreTMX mice to account for deficits in vascular delivery. It also cannot be ruled out that an endurance exercise test may have revealed a performance deficit. A greater reliance on intracellular glycogen and/or lipid stores may sustain short term peak exercise if vascular delivery does not match work rate. EC itgβ1^fl/fl^SCLcreTMX mice have exercise intolerance beginning ∼28 days post-TMX. The increased % body water suggests early stages of tissue edema. This suggests that the accumulation of fluid in the interstitium cannot be properly managed by the lymphatic system. This could increase interstitial pressures and impair delivery of nutrients to working muscle thus contributing to exercise intolerance. EC itgβ1 can sense increases in flow velocity and initiate a signaling cascade resulting in eNOS-mediated signaling in resistance vessels (27, 28). Stimulation of nitric oxide-cGMP signaling results in vascular relaxation and increased insulin-stimulated muscle glucose uptake (53). Interestingly, mice devoid of muscle VEGF-A (54) phenocopy the insulin resistance and capillary rarefaction caused by a decrease in EC itgβ1 in the present study, supporting the idea that EC integrins can regulate VEGF receptor signaling (55).

Capillaries of the microcirculation are the conduit for vascular – parenchymal cell molecular exchange (2, 53). Glucose and insulin are the two most basic determinants of cellular glucose uptake. The rate of capillary insulin efflux is rate limiting for insulin action *in vivo*. A reduced rate of rise in interstitial insulin during an insulin clamp is characteristic of obesity and diabetes (8). The rate that it takes for insulin to reach the skeletal muscle depends on the rate of insulin dispersion across high resistance capillary ECs and the number of capillaries that perfuse the tissue (5, 6). We find that the kinetics of insulin efflux from capillary lumen (e.g. gradient decay constant) is not significantly different in itgβ1^fl/fl^SCLcreTMX and itgβ1^fl/fl^TMX mice. The decrease in muscle capillaries, as defined by CD31+ cells, in EC itgβ1^fl/fl^SCLcreTMX mice is expected to limit access of insulin to the target tissue. We tested whether insulin access was functionally reduced by examining whether activation of insulin signaling proteins was decreased. Muscle insulin signaling was equivalent in response to insulin in itgβ1^fl/fl^SCLcreTMX and itgβ1^fl/fl^TMX mice at the end of the insulin clamp. Tissue glucose uptake is reduced in itgβ1^fl/fl^SCLcre mice despite similar insulin signaling. In addition, [^14^C]2-deoxyglucose uptake is not impaired in skeletal muscle from itgβ1^fl/fl^SCLcreTMX mice compared to itgβ1^fl/fl^TMX mice *ex vivo*. This supports the conclusion that the site of impairment to glucose uptake is the microcirculation, but that this impairment is not due to a deficit in insulin access.

Microcirculatory blood flow velocity, capillary density, and blood glucose determines the perfusion-limited delivery of glucose (41, 56). After accounting for differences in body water, CD31+ staining, and microcirculatory blood flow we estimate that glucose delivery is reduced by ∼50% in itgβ1^fl/fl^SCLcreTMX mice. The calculated estimate of glucose delivery is supported by the ratio of arterial to tissue glucose concentration. This ratio can be reduced because glucose fractional extraction is increased or because glucose delivery is impaired. The former can be ruled out because glucose extraction is impaired during insulin stimulation in itgβ1^fl/fl^SCLcreTMX mice. These results support the concept that impaired glucose delivery is the basis of the reduction in skeletal muscle glucose uptake of itgβ1^fl/fl^SCLcreTMX mice. The present study was designed to focus on skeletal muscle because of their higher rate of glucose uptake and its larger contribution to body mass. Collectively, studies in skeletal muscle support the tenet that EC itgβ1 receptors exert a greater influence on perfusion-limited molecules (e.g. glucose) than on insulin, which is mainly dispersion-limited.

In summary, we developed a mouse itgβ1 knockdown model to study the contribution of EC itgβ1 *in vivo* without compromising health status, based on physiological criteria. We show that 30% reduction in EC integrin β1 is sufficient to cause microcirculatory dysfunction and lead to insulin resistance. This study emphasizes the importance of EC itgβ1 on microcirculatory function and the importance of microcirculatory function on the ability of muscle to consume glucose.

## Acknowledgments

We acknowledge the following Vanderbilt University (VU) and Vanderbilt University Medical Center (VUMC) core facilities: VUMC Hormone Assay & Analytical Services Core (NIH DK135073 and DK020593), VU Metabolic Mouse Phenotyping Center [VMMPC (NIH DK135073; www.vmmpc.org)], and Translational Pathology Shared Resource (NCI/NIH Cancer Center Support Grant 5P30 CA68485-19). We thank Tasneem Ansari, Staci Bordash, Teri Doss, Alicia Kellarakos, Carlo Malabanan for their assistance with surgical procedures and clamp studies. We also thank Dr. Dawn Newcomb, Peter Gulleman, and Dr. Alice Hackett for technical assistance.

This work is dedicated to Dr. David Wasserman, a great scientist, colleague, and mentor.

